# Questioning the G2 phase in the budding yeast cell cycle with a qualitative and possibilistic model

**DOI:** 10.64898/2026.02.06.704310

**Authors:** Adrien Fauré, Dimitris Liakopoulos, Cédric Gaucherel

## Abstract

The budding yeast *S. cerevisiae*, a foundational model for cell cycle studies, exhibits a complex phase organisation (G1, S, G2/M) governed by checkpoints ensuring faithful cellular inheritance. However, the existence of a distinct G2 phase in yeast remains debated, with some advocating for a prometaphase instead. To address this issue, we developed a discrete-event, qualitative, and possibilistic model, the first one to our knowledge, to integrate organelle-level components (replication forks, sister chromatids, mitotic spindle, bud) while remaining parsimonious. Unlike molecular-centred or overly complex whole-cell models, this approach bridges broad systemic and finer mechanistic scales. Our results demonstrate that the model faithfully recapitulates cell cycle progression and supports the dispensable G2 phase. This possibilistic model inspired from recent applications in ecology advocates in favor of the necessity of prometaphase. This study thus provides a unifying and flexible framework to resolve long-standing ambiguities in yeast cell dynamics, while avoiding the pitfalls of excessive complexity or reductionism.

## Introduction

The budding yeast *S. cerevisiae* has been one of the founding model organisms for the study of cell cycle phases, through the discovery of cell division cycle (*cdc*) mutants (Basu et al., 2022; Hartwell et al., 1970; Uzbekov and Prigent, 2022). In this organism as in other eukaryotes, the succession of the different phases is monitored by various checkpoints, that control the correct timing and completion of each cell cycle event (Hartwell and Weinert, 1989; Nurse, 2000). Indeed, before dividing, an eukaryotic cell must not only replicate its DNA and segregate the replicated chromosomes, but also ensure that each daughter cell inherits a full set of cellular components, a process which involves the complex and dynamic assembly and disassembly of specialized structures and compartments, such as the mitotic spindle (Liakopoulos, 2021) and the nucleus (Santana-Sosa et al., 2023).

Typically in eukaryotic cells, phase S of the division cycle corresponds to the period during which DNA is replicated, and phase M, to the different stages of mitosis: spindle assembly, kinetochore attachment, spindle elongation, ending with cytokinesis or cell division proper (Hartwell, 1974; Nurse, 2000). Phase G1 occurs between M and S, and phase G2, between S and M. In contrast with other eukaryotes, however, budding yeast DNA replication and spindle formation can occur simultaneously, as blocking the onset or the completion of S phase does not impair the formation of a mitotic spindle. Therefore, although a G2 phase has been described in yeast cells (Pizzul et al., 2022), its existence is controversial, and a prometaphase has been proposed instead (Tanaka et al., 2005).

Modeling is an appropriate tool to address this issue, which requires an integrated understanding of the cell as a whole, including processes such as DNA replication, spindle pole body duplication, spindle formation, chromosome separation, budding or indeed cytokinesis itself. Surprisingly few studies have tackled this challenge. While early models addressed the connection between cell cycle duration, cell size and population growth (e.g., Basu et al., 2022; Csikász-Nagy and Mura, 2016), in the wake of molecular cell biology, the focus has shifted to the study of a molecular network at the center of which lay the cyclins, their Cdk partners, together with their activators and inhibitors: the so-called *core engine* of the cell cycle (Basu et al., 2022; Nurse, 2000). Some of these models may include cell-level components or processes, such as the mass, bud, spindle and origins of replication (Chen et al., 2000; Williams et al., 2023), but the focus remains on the molecular machinery (Csikász-Nagy and Mura, 2016). Logical models have globally followed the same path as ODE-based models (Csikász-Nagy and Mura, 2016; Fauré and Thieffry, 2009).

Recently, fueled by the progress in the -omics sciences and computational power, *“whole-cell”* models have started considering an integrative approach, incorporating ever more detailed data on metabolites, proteins, nucleic acids, the cell membrane, complete with reaction rates, realistic molecular movement and spatial localization, at different scales (Karr et al., 2012; Thornburg et al., 2022). This latter approach relies on the fact that all the processes occurring during the cell cycle can be simulated from molecular-level interactions. However, it lacks a unifying theory articulating involved processes and a flexible method for handling them. As such, these models are highly complex and heterogeneous, and while they can yield predictions, they hardly explain how cell is functioning as a system (Waltemath et al., 2016).

In this context, we evaluated the feasibility of a discrete, qualitative and possibilistic model (Betz, 2010; Thomas et al., 2022), which could simultaneously represent a whole cell in its various components, while remaining simple enough to be intelligible and exact (i.e., fully controlled). To the best of our knowledge, the model we present here is the first one to apply such methods to an organelle-level description of the cell, while it proves useful in similar integrated challenges in ecosystem ecology (Gaucherel et al., 2024; Pommereau et al., 2022a). The cell cycle of *S. cerevisiae* has already been modeled in a discrete formalism resembling ours (Abou-Jaoudé et al., 2016; Bornholdt, 2008; Fauré et al., 2009), with a more traditional focus on the cyclin / Cdk engine. In contrast, our model focuses on the subcellular scale, incorporating components and processes such as replication forks, sister chromatid cohesion and separation, the spindle pole bodies (SPB) and the mitotic spindle, the bud, as well as cell size.

For the ease of reading, we chose here to gradually present the cell EDEN model, then the analysis of its computed dynamics, and the slightly modified model splitting the G2 focused process, with a gradual discussion on each point. This allows progressively justifying the light changes at each step to improve the cell cycle modeling. We first showed that the model recapitulates the expected succession of events occurring during the cell cycle. To address the existence of a prometaphase (Tanaka et al., 2005) vs. a G2 phase, we then analysed our model from the point of view of these two definitions. Our results show that the model’s dynamics can indeed bypass the G2 phase, whereas the prometaphase appears mandatory. Hence, the present model supports the latter definition (Tanaka et al., 2005), which also seems more consistent with the data.

## Methods

### Background

Our model is designed to account for what happens at the sub-cellular level during the cell cycle of the budding yeast *S. cerevisiae*. The model thus includes variables (Table 1) that represent the cell itself (its size, the presence of a bud, the cytokinetic apparatus that will perform cytokinesis by contracting the bud neck), chromosomes (replication forks, sister chromatids, the cohesin that holds them together, and the presence of two sets of chromosomes towards the end of the cycle), and the mitotic spindle (spindle pole bodies before and after one of them enters the bud, the spindle itself, and kinetochores). Further, we give an overview of the expected sequence of events, along with the cell cycle phases.

**Table 1.** The list of all system components used in the EDEN model, with their associated categories (first column), variable names (second column), and short descriptions (last column).

|  | component | full name & description |
| --- | --- | --- |
| cellular components | siz | cell size |
|  | bud | the bud |
|  | sep | septin ring (Lew, 2003) |
| chromosomes | frk | replication forks; appear after initiation of DNA synthesis |
|  | csn | cohesin; holds sister chromatids together |
|  | chd | sister chromatids, *i.e.* duplicated chromosomes, still attached after S phase is completed. Note that some regions of the chromosomes may still be unreplicated, but the bulk of DNA is. |
|  | ch2 | (fully) replicated chromosomes; double the G1 nuclear DNA content = post replication nuclear DNA |
| spindle | SPB2 | duplicated spindle pole bodies, before one of them enters the bud |
|  | SPBB | spindle pole bodies, duplicated and inserted in the nuclear envelope, after one of them has entered the bud |
|  | spn | mitotic spindle, after SPB separation (Jacobs et al., 1988; Jaspersen and Winey, 2004) |
|  | kin | kinetochores, attached to microtubule, ready for sister chromatid separation |

The yeast cell cycle begins in phase G1 with *start*, an event which marks cell cycle entry but also the commitment to complete an entire cell division (Hartwell, 1974; Hartwell et al., 1974). The exact mechanism behind *start* is still actively discussed (Johnson and Skotheim, 2013). Here we represent it through a simple (*siz*) variable, representing cell size and, indirectly, the balance between the G1 cyclin Cln3 and the G1 transcription inhibitor Whi5 (Facchetti et al., 2017).

Following phase G1, yeast cells enter phase S, where they replicate DNA and initiate bud growth (Hartwell, 1974). At the end of the S phase, cells duplicate the spindle pole body (Jaspersen and Winey, 2004; Matellán and Monje-Casas, 2020), an event that marks the beginning of bipolar spindle assembly. Chromosomes attach to the spindle at the level of the centromere, through a protein complex called the kinetochore (McAinsh and Marston, 2022). In other eukaryotes, this process happens at the beginning of mitosis, after a gap phase called G2. In budding yeast in contrast, kinetochore attachment begins as soon as the centromere has been replicated in S phase, and this difference has led to question the existence of a G2 phase in this organism, and to propose a prometaphase instead (Tanaka et al., 2005).

At the end of yeast metaphase, the bud has grown to 2/3 of the mother size and all kinetochores are bound in a bipolar manner. When this has been achieved, the cell moves into anaphase. Here, cohesin is degraded and the sister chromatids separate to two equal sets that are segregated between mother and bud. Finally, the spindle disassembles, cytokinesis and cell separation follow, resetting the cell to its initial state (Fraschini, 2016; Meitinger and Palani, 2016).

Throughout the cell cycle, molecular mechanisms called *checkpoints* ensure an orderly sequence of events (Hartwell and Weinert, 1989). Hence, the start checkpoint delays the initiation of DNA synthesis until the cell has reached a large enough size (Johnson and Skotheim, 2013). The DNA replication checkpoint protects stalled replication forks and prevents segregation of unreplicated chromosomes (Ciardo et al., 2019). The spindle assembly checkpoint (SAC) prevents sister chromatid separation until all chromosomes are properly attached to the spindle (Hayward et al., 2019). The spindle position checkpoint (SPoC) delays mitosis until one SPB has entered the bud and the spindle is correctly positioned along the mother-daughter axis (Matellán and Monje-Casas, 2020). Drugs or mutations that prevent bud growth lead to the activation of a morphogenesis checkpoint, that prevents delay entry into mitosis until bud growth can resume (Lew, 2003). Other checkpoints mechanism monitor DNA damage (Zhang et al., 2009), membrane integrity (Kono and Ikui, 2017), unsegregated DNA lagging through the bud neck (Mendoza et al., 2009), or stress to the endoplasmic reticulum (Niwa, 2020). Together, these mechanisms contribute to the survival of the cell lineage, which we did not directly modeled instead of let them emerging *a posteriori* from the computed dynamics.

### EDEN modeling framework

For modeling the cell cycle in a whole-cell spirit, we used here models from the large family of discrete-event models (e.g., Chaouiya and Remy, 2013; Reisig, 2013), closely related to Petri nets and Boolean networks (Abou-Jaoudé et al., 2016). We developed a model from the EDEN framework (Gaucherel et al., 2024; Pommereau et al., 2022a), allowing possibilistic modeling as well as a flexible combination of asynchronous and synchronous processes required for such a challenge. Such a model is first defined by a set of Boolean variables, whose values are denoted as + (‘On’ or functionally present) or - (‘Off’, functionally absent), representing the qualitative states of the system components. The whole system state is defined as a vector storing the values of all the variables of the studied system. We used this vector for defining the initial state from which the dynamics will be computed. The model is then defined by a set of “if-then” reaction rules, which describe transitions between system states. Each rule is made up of a left side formalizing its conditions of application, in terms of which variable(s) should be On or Off for the rule to be “fired”. The right side of the rule formalizes the realizations and expected state changes, in terms of the set of variables that will be switched to On or Off (or possibly, left unchanged).

Regarding rule definition, two important features stand out in EDEN models. In a traditional discrete-event formalism (e.g., Abou-Jaoudé et al., 2016; Reisig, 2013), logical functions usually define the target level for each variable, and the update scheme (most commonly synchronous or asynchronous, though several others exist) is applied *a posteriori*. Conversely, in the EDEN formalism, variables are stable by default, and only variables susceptible to change are listed in right sides of the reaction rules. Moreover, while rules are treated asynchronously, when several variables are present on the right side of a rule, the corresponding changes are carried on synchronously (Gaucherel et al., 2024; Pommereau et al., 2022a). Hence, contrary to other Boolean frameworks, the update scheme that determines the dynamics of the system and is usually seen as distinct from the model, is here embedded within the model itself, and may combine features of the synchronous and asynchronous updates.

Together, variables and rules contribute to build two important graphs and associated topologies. Firstly, a *regulatory graph* (also called interaction network or influence network (Chaouiya and Remy, 2013; Comet et al., 2013; Gaucherel et al., 2024; Naldi et al., 2018), whose nodes represent the variables, and whose edges denote the existence of at least one reaction rule, such that for a given edge, the variable represented by its foot appears in the left side (conditions) of the rule, and the variables represented by its head appears on the right side of the same rule. Secondly, a *state transition graph* (STG, also called the dynamics), whose nodes represent distinct states of the system, and whose edges represent transition between these states, as computed by the reaction rules from the chosen initial state. This STG is computed once and delivers the possibilistic dynamics of the modeled system (Gaucherel et al., 2024; Pommereau et al., 2022b). We recall here that a possibilistic modeling framework, such as EDEN used here, is by definition non-deterministic (*i*.*e*. several outcomes from each state) and non-probabilistic (*i*.*e*. no probability or any other weight on transitions), which is a striking difference with traditional modeling frameworks in living sciences.

For the purpose of analysis or simply readability, computed states and transitions may be grouped according to topological or user-defined properties, yielding graphs whose nodes represent set of states that share the same relevant properties (Pommereau et al., 2022a; Thomas et al., 2022). In particular, this study will further exhibit “phase graphs” which are merged STG separating such state sets and their transitions (or checkpoints) in-between, according to pre-defined definitions of each cell phase. Indeed, checkpoints are partly arbitrary chosen, and are defined *a posteriori* as transitions between each cell phase. For this reason, they correspond to switch of specific combinations of the available model variables.

## Results and Discussion

### The computed dynamics

The model was built iteratively, through a trial and error procedure, and seems to be an appropriate trade-off between a simplistic and an over-complicated cell model. We selected 11 variables for this theoretical and parsimonious model, grouped into three main categories (Table 1):

- variables denoting cell growth: cell size (*siz*), the bud (*bud*) and the hourglass-shaped septin collar (*sep*) that forms at the bud neck and monitors bud growth.
- variables denoting DNA at different stages of the cycle: replication forks (*frk*), sister chromatids (*chd*), cohesin (*csn*) that holds sister chromatids together, and finally, fully replicated chromosomes (*ch2*)
- variables denoting spindle components: duplicated and separate spindle pole bodies (*SPB2*), the spindle itself (*spn*) after SPB separation, kinetochores all properly attached to the spindle (*kin*), and the presence of one SPB in the bud (*SPBB*).

Reaction rules were written to translate known biological connections between these components, i.e., processes involving them (Table 2). For example, the *start* checkpoint signals the cell to initiate DNA synthesis, bud growth and SPB separation. Thus, the variable *siz* appears as a necessary condition in the rules activating *frk* (replication forks), *bud* and *SPB2* (spindle pole body duplication). Other checkpoints translate similarly to reaction rules: the SAC implies that attached kinetochores *kin+* and chromatid duplication *chd+* are necessary conditions for cohesin deposition *csn-*; the SPoC, that SPB segregation into the bud *SPBB+* is necessary for complete chromosome segregation (*chd-, ch2+*); and the morphogenesis checkpoint implies that bud growth, denoted by the deformation of the septin ring into an hourglass-shaped collar (*sep+*) is a condition for kinetochore attachment (*kin+*).

**Table 2.** The list of all system processes used in the EDEN model, with their associated reaction rules and short names (first column), and their formal writings (second column). See supplementary materials and Table 6 for details and references.

| name | rule |
| --- | --- |
| [start] | siz- >> siz+ |
| [frk+] | siz+, chd-, ch2- >> frk+ |
| [chd+, csn+] | chd-, frk+ >> chd+, csn+ |
| [SPB2+] | siz+, SPBB- >> SPB2+ |
| [spn+] | SPB2+ >> spn+ |
| [kin+] | sep+, chd+, spn+ >> kin+ |
| [bud+] | siz+ >> bud+ |
| [sep+] | siz+, bud+ >> sep+ |
| [SPB2-, SPBB+] | bud+, SPB2+, chd+, csn- >> SPB2-, SPBB+ |
| [csn-] | kin+, chd+ >> csn- |
| [frk-, chd-, ch2+] | frk+, chd+, csn-, sep+, kin+, SPB2-, SPBB+ >> frk-, chd-, ch2+ |
| [kin-, spn-] | ch2+ >> kin-, spn- |
| [CK] | bud+, sep+, SPBB+, kin-, spn-, ch2+ >> bud-, sep-, siz-, SPBB-, ch2- |

To a large extent, reaction rules depend on the chosen level of details. Some rules affect several variables: this is typically the case of cytokinesis that resets several variables to their initial states. Sometimes however, the interaction is more subtle, as some variables represent different facets of the same object/phenomenon. For example, open replication forks (*frk+*) is a marker of ongoing replication, leading to the formation of sister chromatids, which translates into a reaction rule with *frk+* on the left side, and *chd+* on the right side. Cohesin is laid on DNA during replication, so *csn+* appears on the right side of the same rule. The variables *frk, chd* and *csn* are connected by a causal relationship. In contrast, while the closure of all replication forks marks the end of DNA synthesis, and the switch from sister chromatids (*chd+*) to separate chromosomes (*ch2+*), it cannot be said that replication forks closure *causes* this switch. Rather, these are different ways to describe the same phenomenon, happening all at once when the conditions are satisfied (i.e., one SPB has entered the bud, cohesin have been degraded, and so on). Hence, in the model *frk-, chd-* and *ch2+* are listed on the right side of the same rule, with *SPBB+, csn-* and other conditions on the left.

Finally, some aspects of the model may be arbitrary and other choices could arguably have been made. For example, drawing a parallel with *chd* and *ch2*, we chose to couple together the variables *SPB2* and *SPBB*, which represent the presence of a second spindle pole body, and the entry of a SPBB into the bud, on the right side of the same rule, so that *SPB2* goes down when *SPBB* goes up (Table 2). For the sake of clarity, detailed narratives and bibliographical references backing theoe rules can be found in supplementary materials (Table 6, Supp. Mat.). The cell interaction network is displayed on the basis of these defined reaction rules (Figure 1a). This graph is even what we call a hyper-network in a previous study (Gaucherel et al., 2024) for highlighting the hyper-graph indeed connecting several variables (nodes) to several rules (edges, Figure 1b). This (hyper-)network is quite densely interconnected, reflecting the high degree of coordination of cell division events, and is easily segregating the three main component categories according to their interacting processes.

**Figure 1.**
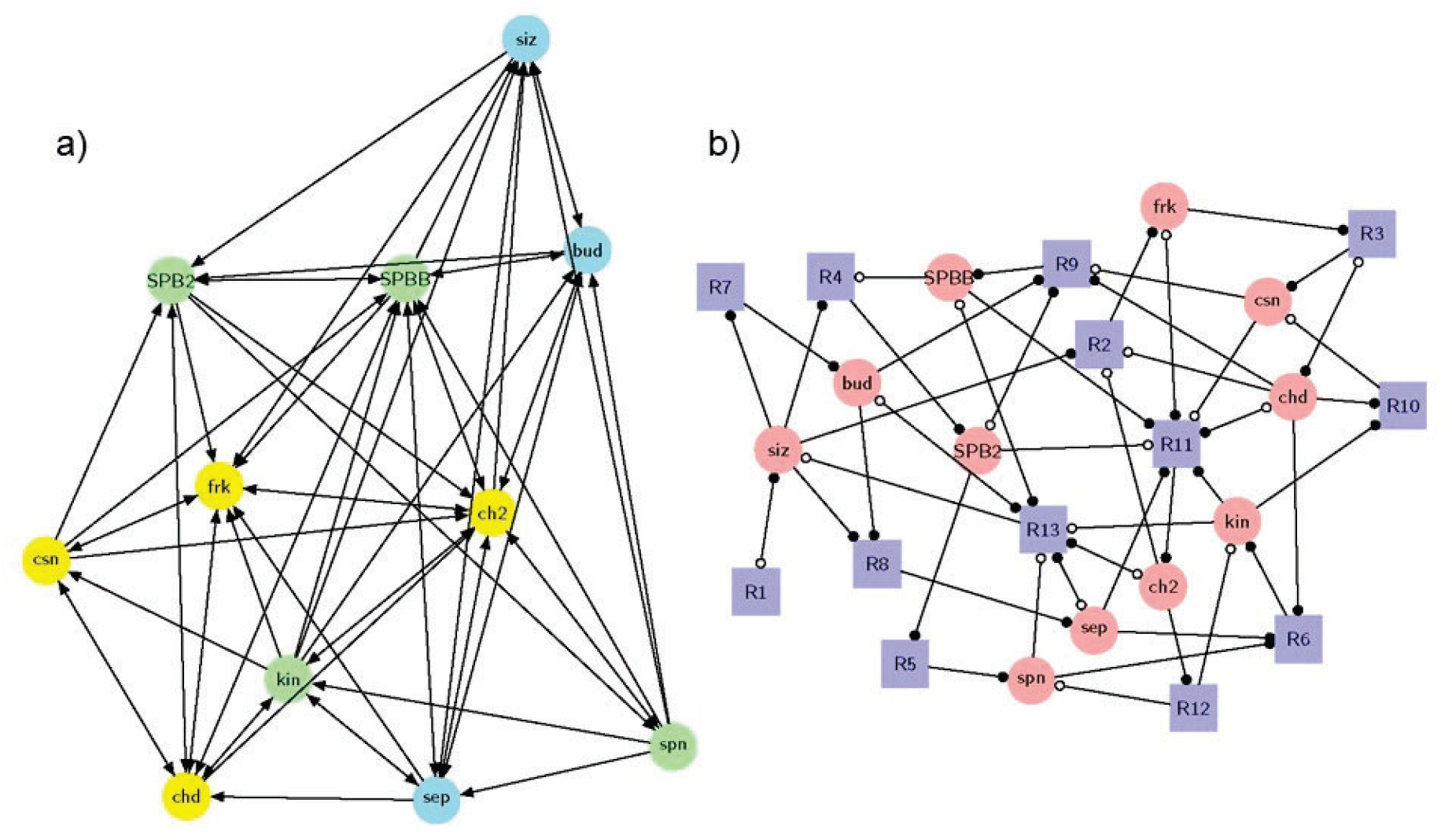
Two displays of the same interaction network for the modeled budding yeast. Depending on the display algorithm used, it can be flattened as a graph(a) or drawn as a more rigorous hyper-graph (b). Colors of the graph (a) denote the three main categories of variables and components, and are the result of the (manual) rule definitions: DNA-related variables in yellow; spindle-related variables in green; cell growth-related variables in blue. Each edge may concern distinct and possibly several reaction rules (*i*.*e*. processes, Table 2). Colors and shapes of the hyper-graph (b) components refer to the type of nodes, being either variables (pink disks, for so-called places) or rules (blue squares, for so-called transitions or rules), while the arc heads show plain (resp. empty) circles for representing positive (resp. negative) variable states in the associated rules (squares). Such round-shape arc heads are pointing squares (resp. disks) for defining rule conditions (resp. rule realisations). See (Gaucherel et al., 2024; Pommereau et al., 2022a) for details.

### The cell cycle phases

Starting from an initial state with every variable switched off, and applying all possible reaction rules yields a small structural stability (or qualitative attractor) with only 33 states (STG nodes), where all variables oscillate (Figure 2). No sub-region (specific topologies) of the computed dynamics may be identified in this stability, which is also called a strongly connected component (SCC, e.g., Pommereau et al., 2022a; Reisig, 2013). By definition, a SCC is a set of mutually reachable states of the system.

**Figure 2.**
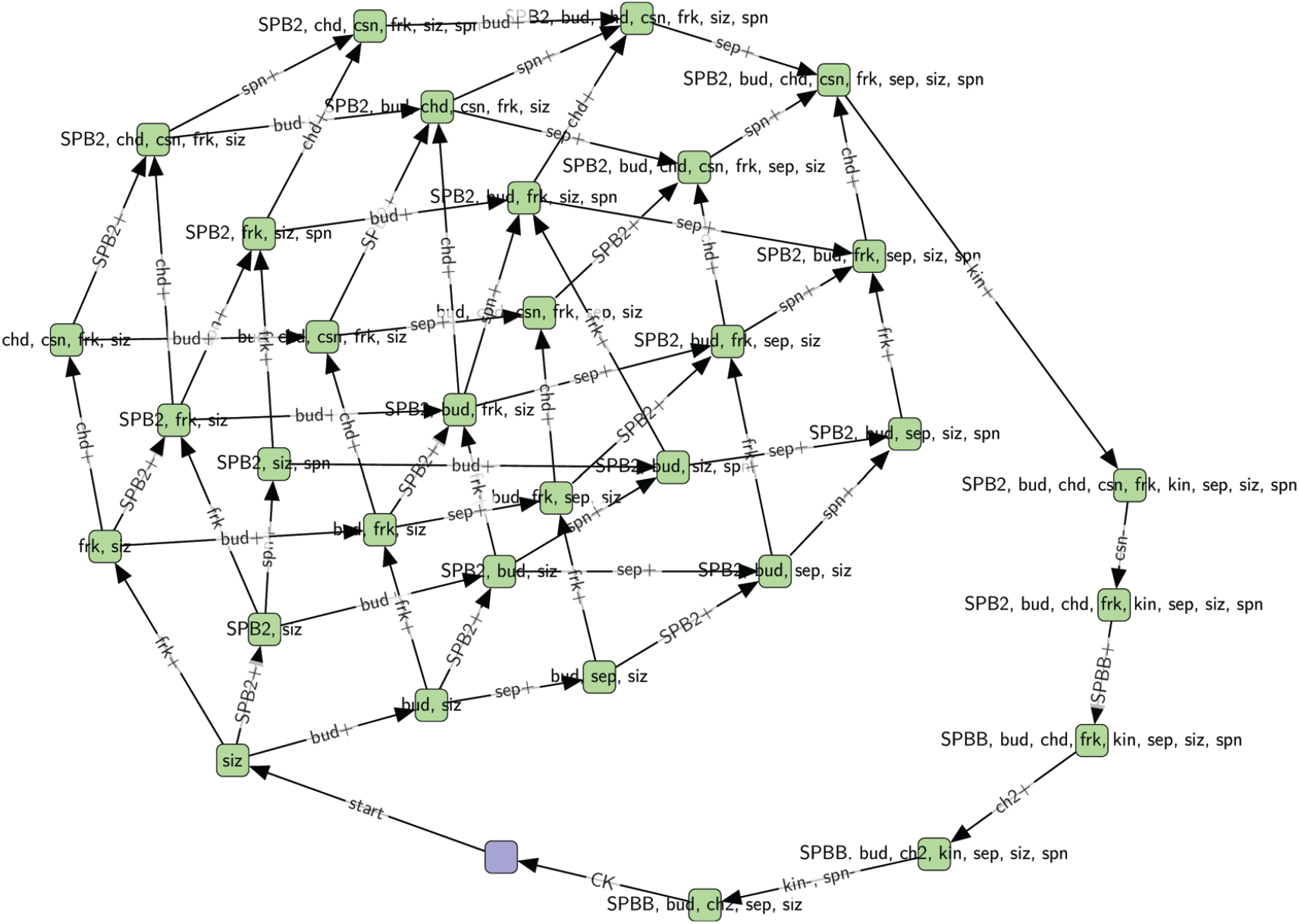
The full computed dynamics (STG) for the wild-type cell. Each node represents here a distinct state of the system, with node labels denoting which variable is On in each state (absent variables are thus switched Off). Edge labels denote which rule fires along to the corresponding transition, with its direction highlighted. The initial state is highlighted in blue. The graph display tends to locate closer system states that are connected through single transitions.

Starting from the initial state, the cell must first reach a critical size (*start* checkpoint), after which three processes start in parallel (right after the *start* transition in Figure 2):

- DNA replication: opening of the replication forks (*frk+*, rule n°2, Table 2), and formation of sister chromatids (*chd+*, rule n°3);
- spindle formation: duplication and separation of the SPB, (*SPB2+*, rule n°4), followed by spindle assembly (*spn+*, rule n°5).
- budding (*bud+*, rule n°7), up to the deformation of the septin ring into an hourglass-shaped collar (*sep+*, rule n°8).

The dynamics then generates multiples pathways which converge to a single state where all the three processes have been completed. Then, the dynamics follows a single linear trajectory, with the bipolar attachment of the kinetochores to the spindle (*kin+*), the degradation of cohesin (*csn-*), followed by the migration of one of the SPB into the bud (*SPBB+*) and completed separation of the sister chromatids into two fully separate chromosomes (*chd-, ch2+*). Then the spindle disassembles (*spn-, kin-*) and the cell exits mitosis and splits, resetting the dynamics to its initial state.

Hence, the dynamics of the attractor matches the expected sequence of events (Basu et al., 2022; Hartwell et al., 1970; Uzbekov and Prigent, 2022). Remarkably, although we observe parallel pathways, showing that some events can be completed in different orders (a strength of the present model), we do not detect any shortcut or any smaller-scale oscillation within this main stability. This is fully consistent with the knowledge that the cell cycle must be completed before a new cell cycle starts. In the next section, we focused on the possible division of this reference dynamics into various cell cycle phases.

### Cell cycle phases in the model

In order to assign each state to a particular phase of the cell cycle, we need to formalize phase definitions with respect to the modeled variables. In practice, we can specify a logical formula for each cell phase, which defines a sub-region of the computed dynamics. In this article, we thus consider two definitions of the cell cycle phases.

The first definition divides the cycle into two main phases, S (for synthesis) and M (for mitosis, including metaphase, anaphase and telophase), separated by gap phases G1 and G2 (Table 3, Hartwell, 1974; Nurse, 2000). Although DNA synthesis gives its name to S phase, this is also the period during which spindle pole bodies are replicated, and a bud forms and grows. We thus define S phase as the period during which these processes occur. When all the three are completed, the cell enters G2. Then, phase M is further divided into metaphase, anaphase and telophase, based on the bipolar attachments of kinetochores to the spindle, sister chromatid separation and spindle disassembly before mitotic exit (Pizzul et al., 2022). These arguments provide formal definitions of the phases based on the chosen model’s variables (Table 3).

**Table 3.**
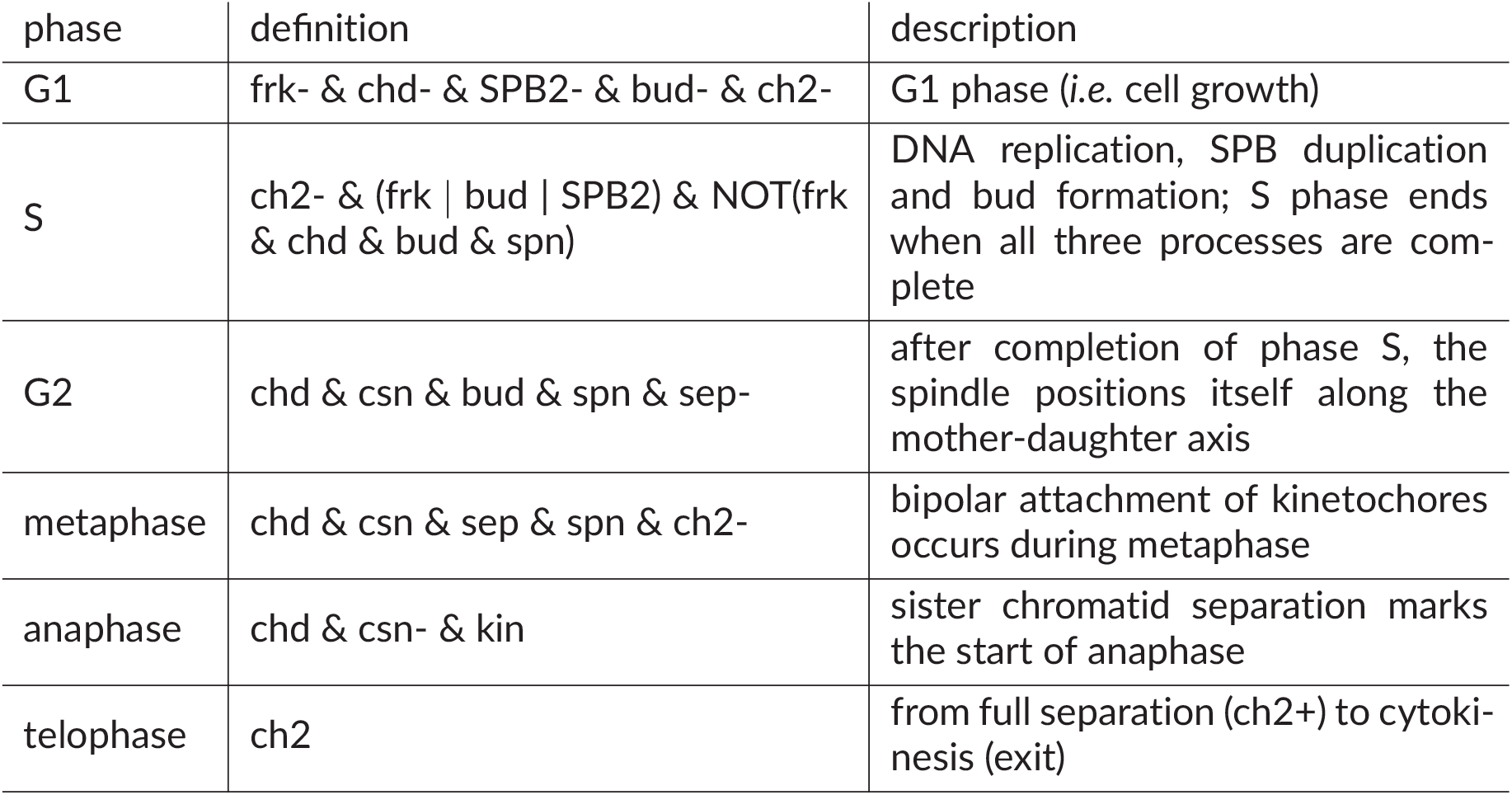
The traditional phase definitions collected from (Hartwell, 1974; Nurse, 2000; Pizzul et al., 2022). The phase names (first column), their formal definitions (second column), and their short descriptions (last column) are given.

The second definition considered contests the existence of a G2 phase, as well as an explicit prophase, in budding yeast (Tanaka et al., 2005). According to Tanaka 2005, we propose that M phase begins with prometaphase, during which microtubules interact with the kinetochores. In our system (Table 4), this translates to the activation of the *chd* and *spn* variables. Then, bipolar kinetochore attachment (*kin+*) moves the cell into metaphase. In contrast to the previous definition, here bipolar attachment thus marks the beginning of metaphase. The other phases remain defined as previously (Hartwell, 1974; Nurse, 2000; Pizzul et al., 2022).

**Table 4.**
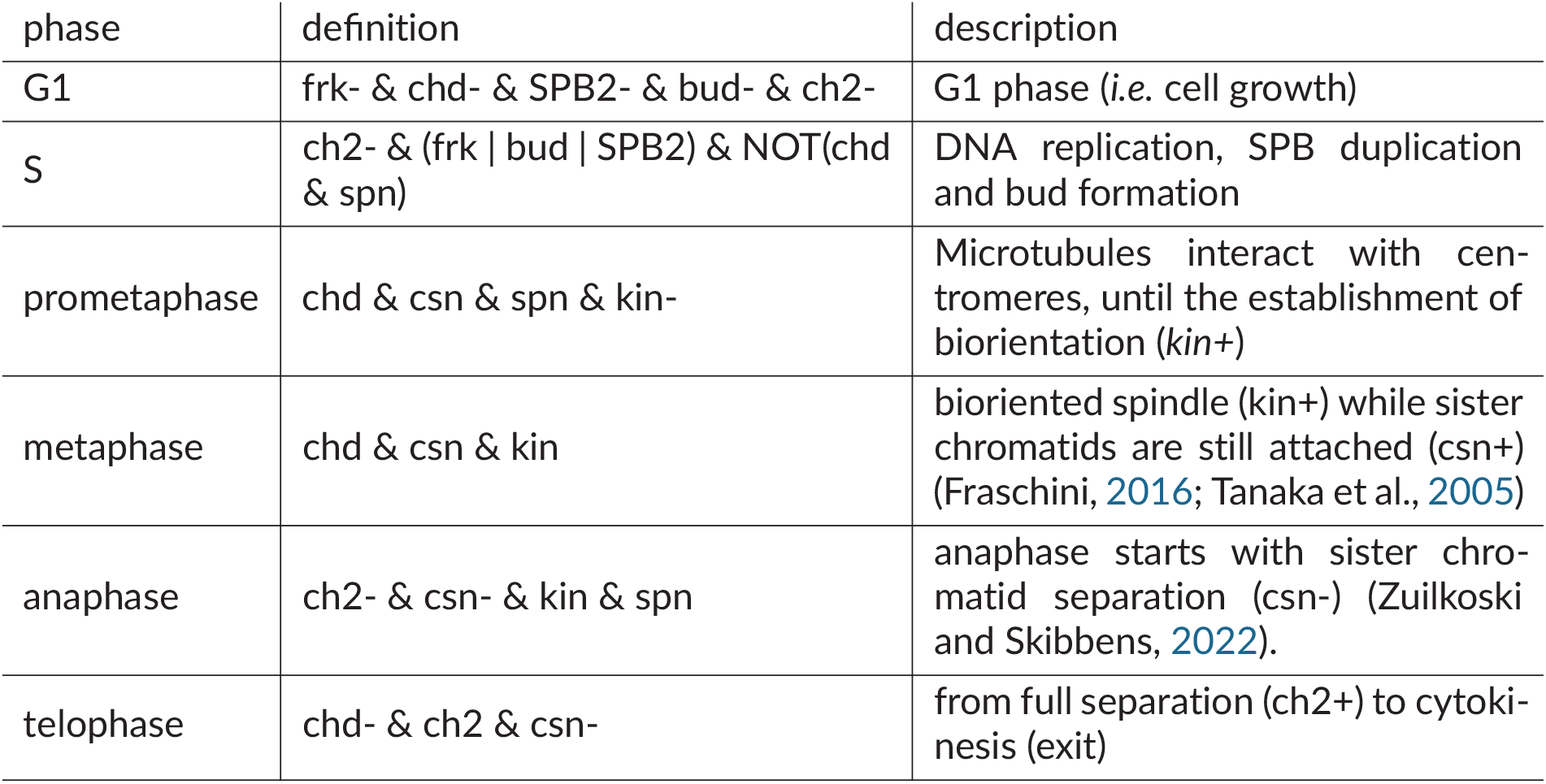
The phase definitions according to Tanaka *et al*., 2005 (Tanaka et al., 2005). The phase names (first column), their formal definitions (second column), and their short descriptions (last column) are given.

### Does the budding yeast cell cycle have a G2 phase?

We first used the traditional phase definition (Hartwell, 1974; Nurse, 2000), which divides as said the cell cycle into two main phases separated by two gap phases (Table 3). Based on this definition, the initial state and its immediate successor belong to G1, until the opening of the replication forks (*frk+*), budding (*bud+* or the duplication of the *SPB2+* move the system to S. Most states in the stability belong to this phase (24 states), the ones produced by the distinct pathways. When all the three starting processes are completed, the cell enters a single G2 state, or directly into metaphase if budding has progressed enough (*sep+*). Surprisingly, this definition leads to the observation that, at least in our model, the G2 phase is optional.

The two metaphase states are then directly followed by anaphase (two states) and telophase (two states), before returning back to the initial G1 state. It is possible to automatically draw such a merged sequence into a phase graph (Figure 3), while the details of each phase content can be found in supplementary materials (Figure 6, Supp. Mat.).

**Figure 3.**
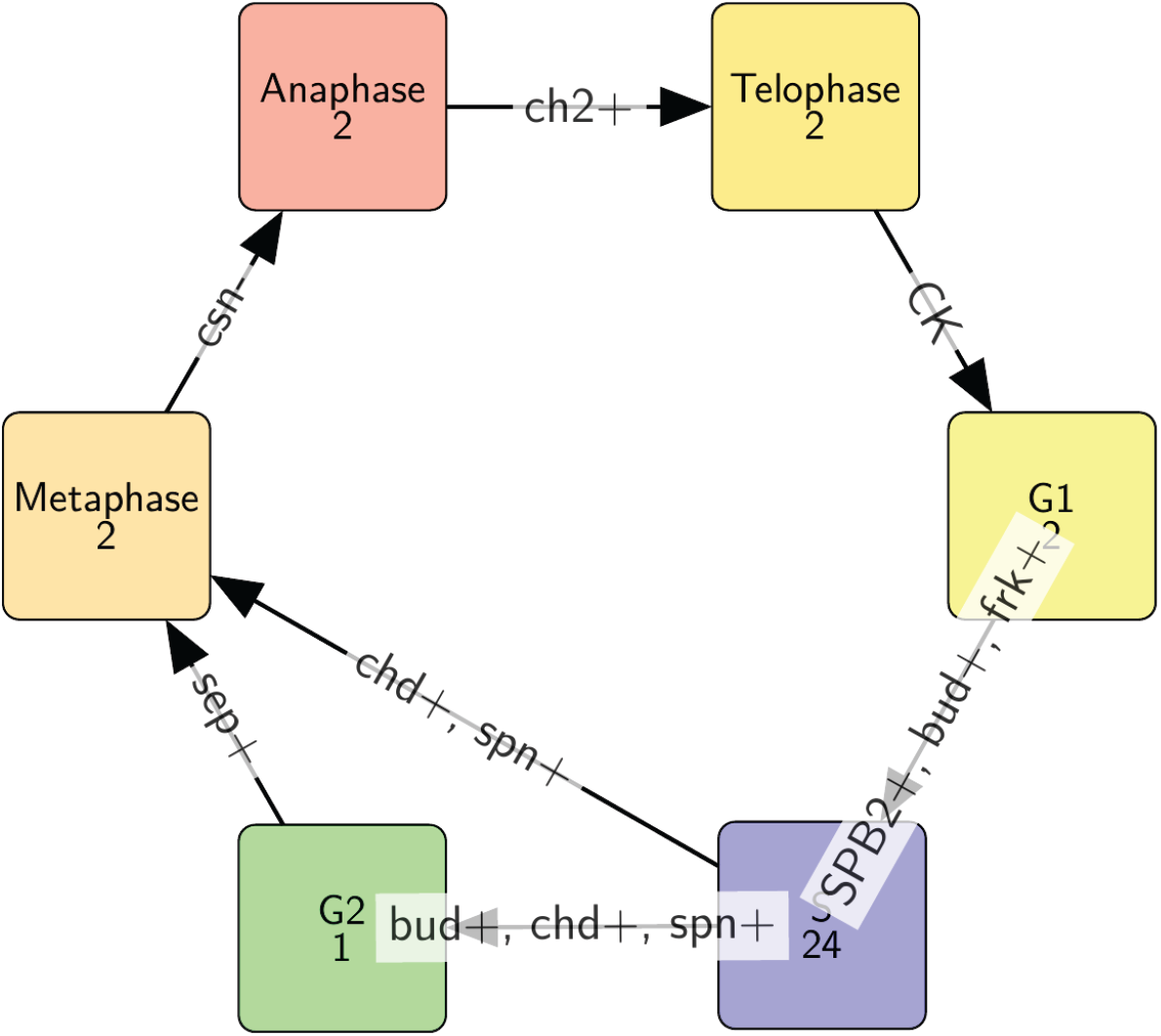
The phase graph of the wild-type cell in which the states are merged according to the phase definitions given in Table 3. Each node of this graph is a set of system states computed from the chosen initial state and using the predefined reaction rules; the labels of the nodes are their names and the number of merged states. The edges are highlighting the transitions between these phases, with their directions and labels indicating which variables should be changed for reaching the next phase.

The second definition contests the existence of a G2 phase, as well as an explicit prophase, in budding yeast (Tanaka et al., 2005). Instead, the authors propose that M phase begin with prometaphase, during which microtubules interact with the kinetochores. In our modeled system (Table 4), this translates as the activation of the *chd* and *spn* variables. Then, bipolar kinetochore attachment (*kin+*) moves the cell into metaphase. In contrast to the previous definition, here bipolar attachment thus marks the beginning of metaphase. The other phases are defined as previously (Table 3). This new definition yields a phase graph quite similar to the previous one, with the initial state and its immediate successor both in G1, followed by distinct pathways opening into a large (23 states) S phase. The difference appears towards the end of phase S, as all states now converge to three prometaphase states (Figure 4), followed by a single metaphase state. Anaphase and telophase are unaffected by the change, while the details of each phase content can be found in supplementary materials (Figure 5, Supp. Mat.).

**Figure 4.**
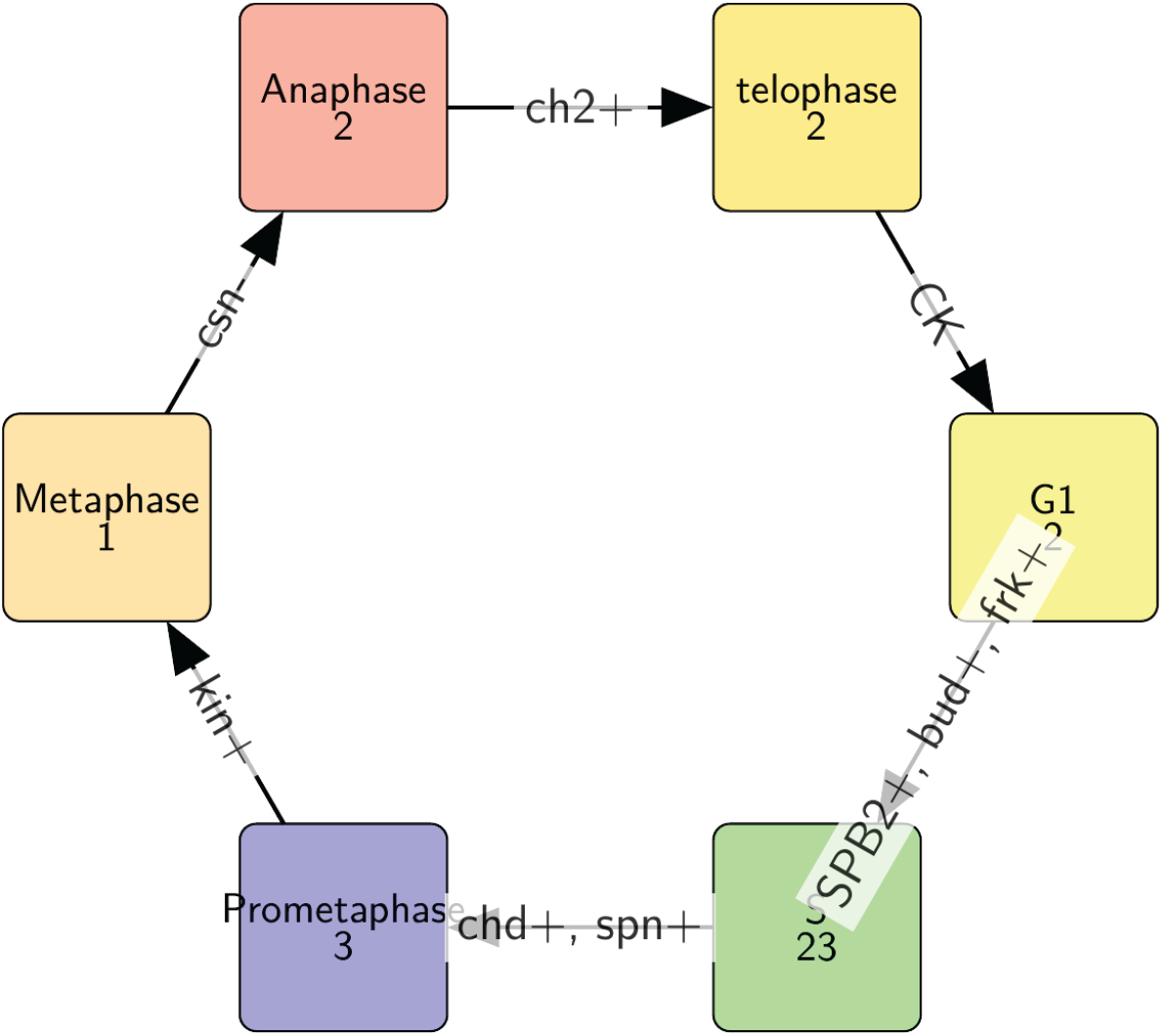
The phase graph of the wild-type cell in which the states are merged according to the phase definitions given in Table 4. Each node of this graph is a set of system states computed from the chosen initial state and using the predefined reaction rules; the labels of the nodes are their names and the number of merged states. The edges are highlighting the transitions between these phases, with their directions and labels indicating which variables should be changed for reaching the next phase.

**Figure 5.**
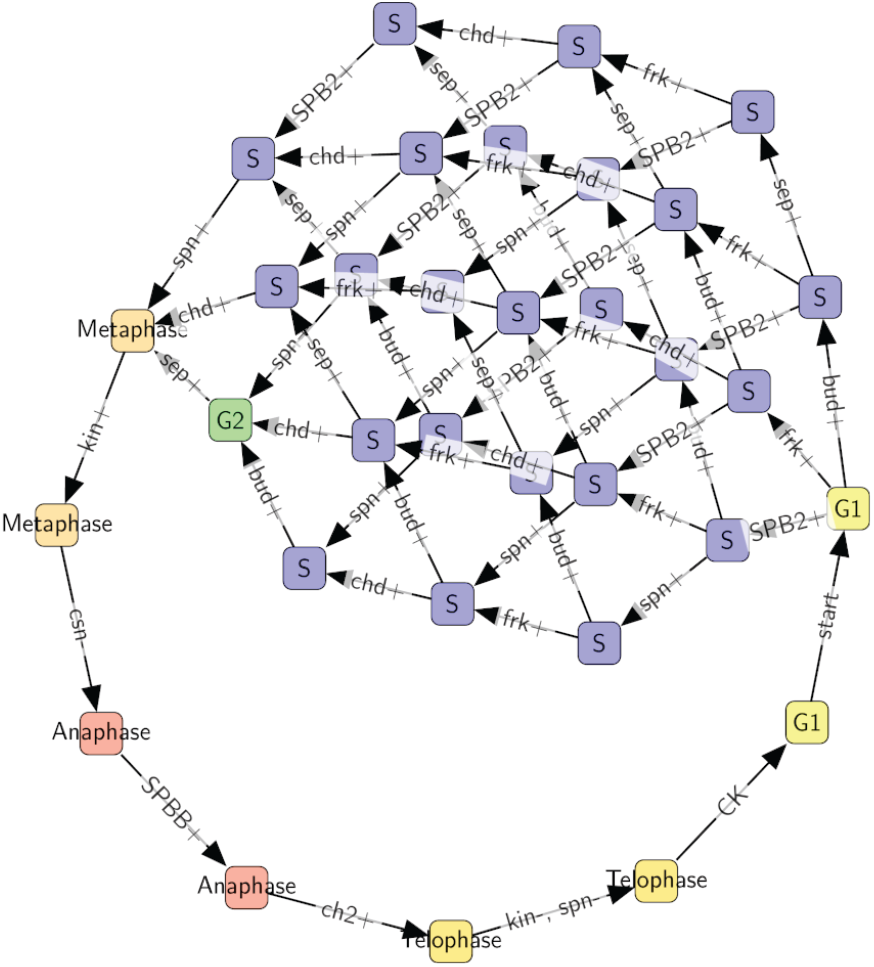
Full state transition graph for the wild-type case (corresponding to Figure 3). States are colored according to the classical phase definition.

**Figure 6.**
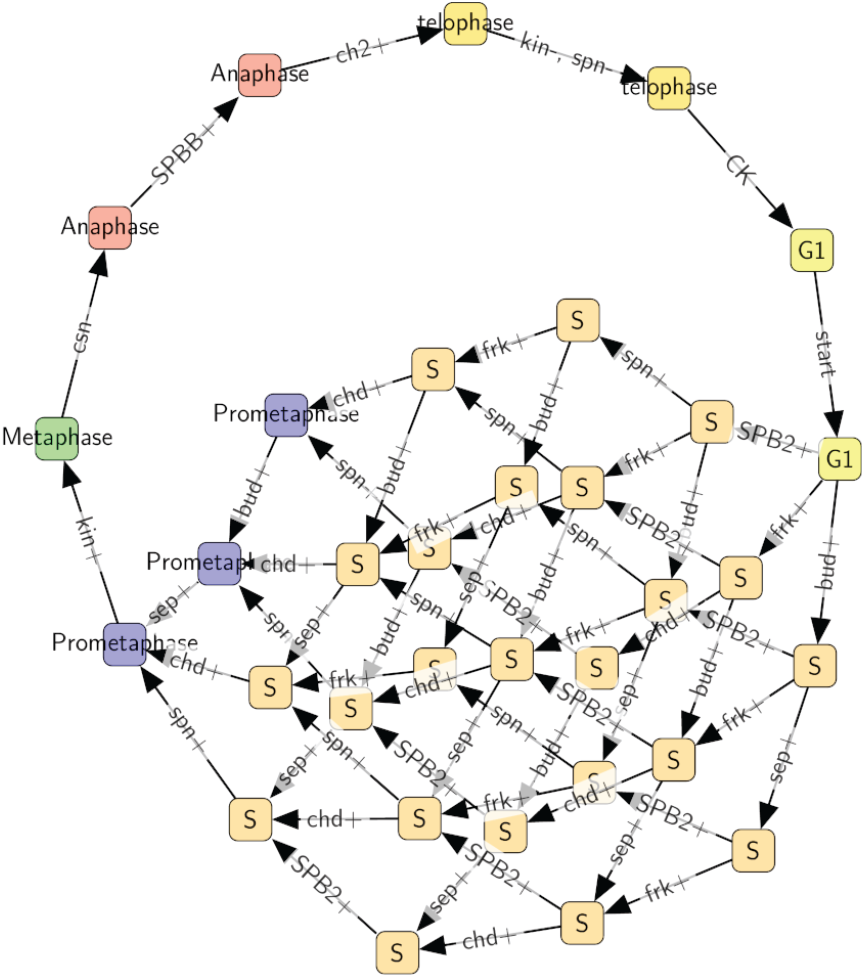
Full state transition graph for the wild-type case (corresponding to Figure 4). States are colored and marked according to the phase definition proposed by Tanaka *et. al*, according to which there is no G2 phase.

### Synchronous reaction rules steer the dynamics

A characteristic of the EDEN formalism as compared to many other discrete-event models, is the ability to write reaction rules with several variables on the right side (the possible realizations or changes), embedding synchronous transitions within the model itself. This places the EDEN formalism within the family of qualitative, set-rewriting-like formalisms. With that in mind, we examined the influence of such rules on the dynamics of the model. The present model shows five such rules, coupling two or more variables in synchronous transitions (Table 2). To evaluate the importance of such couplings onto the dynamics, we run a modified version of the model, in which each of these rules had been broken down into separate rules with the same conditions (left side of the rules), and only one variable as a realization (right side), thus decoupling these transitions. For example, the rule named *[frk-, chd-, ch2+]* is replaced by three rules as follows:

- *[frk-] frk+, chd+, csn-, sep+, kin+, SPB2-, SPBB+ >> frk-*
- *[chd-] frk+, chd+, csn-, sep+, kin+, SPB2-, SPBB+ >> chd-*
- *[ch2+] frk+, chd+, csn-, sep+, kin+, SPB2-, SPBB+ >> ch2+*

With this new model splitting most of these rules (Table 5), the computed dynamics leads to abnormal cycles or deadlocks (i.e., some states from which the system cannot leave, according to the predefined rules). The only exception is the rule *[kin-, spn-]*, whose splitting only replaces the one synchronous transition with two diamond-shaped (concurrent) pathways, after which the cycle resumes unaffected. This is due to the fact that both variables affected by the rules appear in the conditions for the rules applied next within the cycle (*[CK]*), and that no other rule can apply unless both variables have changed. We insist here that the fact to split some rules (realizations) and compute more system states is technically trivial; the main point here are the biological justifications for such split (resp. merging) which are provided in Table 5.

**Table 5.** The decoupling synchronous transitions. This table lists the reaction rules of variants of the first model where single rules affecting multiple variables have been replaced by as many rules affecting each variable independently. The rules are listed here with their short names (first column), and their associated phenotypes (second column) and explanations (last column). See Supp. Mat. Table 6 for detailed narratives and references.

| Rule | phenotype | explanation |
| --- | --- | --- |
| [chd+, csn+] | normal cycle or S phase arrest | activation of *chd* before *csn* leads to premature activation of the [SPB2-, SPBB+] rule; inactivation of *SPB2* prevents the activation of the [spn+] rule, thus blocking cell cycle progression. |
| [SPB2-, SPBB+] | normal cycle, plus *SPB2* oscillations in anaphase | conditions for the [SPB2+] are still valid in anaphase, so inactivation of *SPB2* allows it to immediately activate again. |
| [frk-, chd-, ch2+] | normal cycle, abnormal cycles, or arrest in S phase | decoupling may lead to cytokinesis before the *chd* variable has turned off. This in turn prevents the activation of the [chd+, csn+] rule, which allows for the premature inactivation of *SPB2*, preventing activation of *spn*. |
| [kin-, spn-] | normal cycle | Both variables affected by the rule appear in the condition of the [CK] rule that fires next in a normal cycle, and no other rule can fire unless they both change. |
| [CK] | cell cycle arrest | Most variables affected by the rule are also present in the rule's condition, so if only one of those variables change, the condition for changing the other becomes invalid, and the system reaches a deadlock. |

**Table 6:**
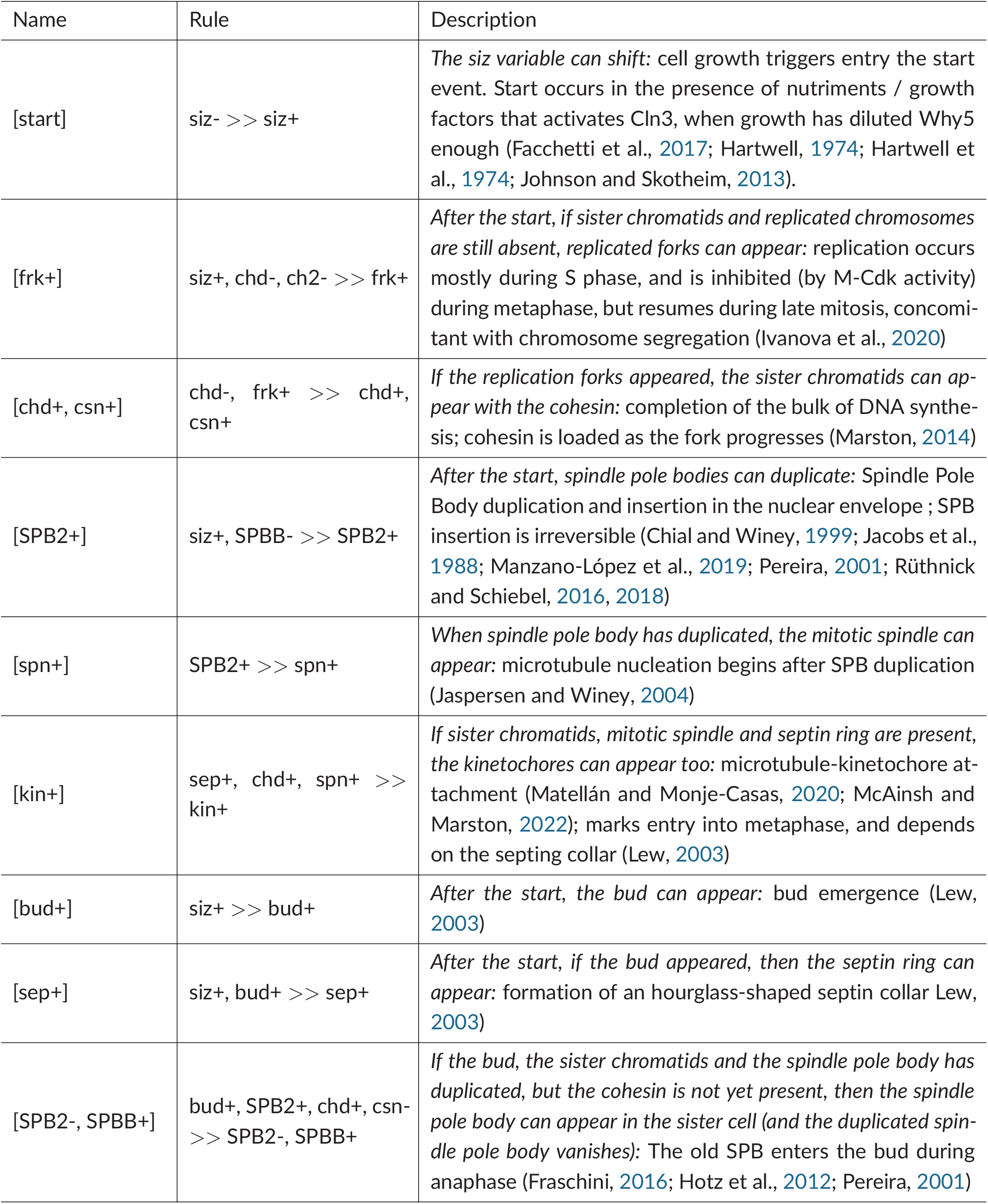
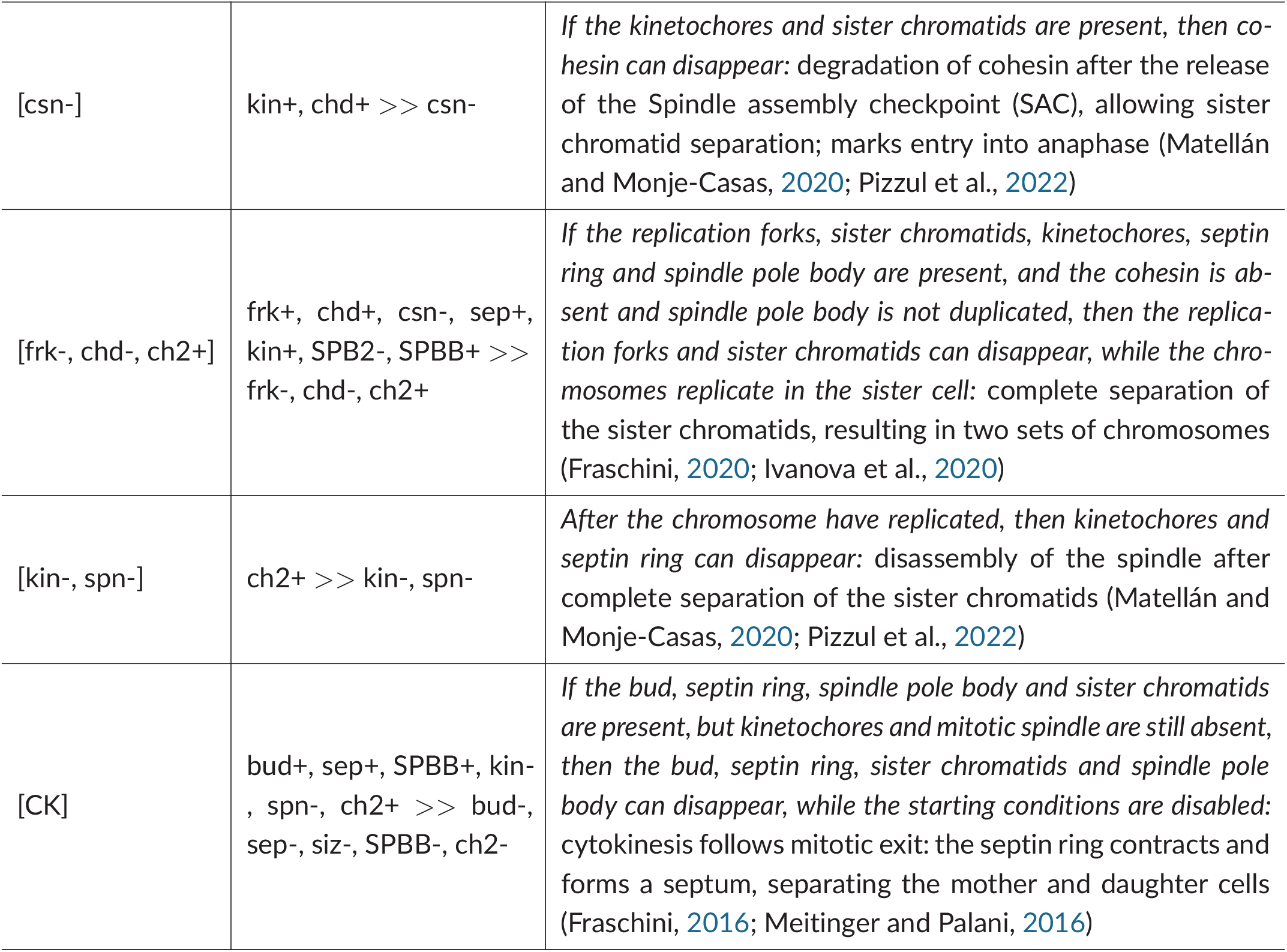
Details of the rules for the EDEN model of this study (see also Table 2). The name (brackets), formal syntax and short descriptions (raw narrative: followed by its interpretation) of each possibilistic rule is listed, and is duly referenced (see references following the main text).

To some extent, these abnormal phenotypes could be salvaged, or aggravated, by adding or removing variables from the rule conditions, in a way that does not affect the dynamics. For example, removing *spn-*from the condition of the *[CK]* rule, does not change the wild-type behaviour; but then splitting the *[kin-, spn-]* rule yields a new trajectory forming a cycle where the *spn* variable remains always On. Similarly, including both *SPB2+* and *SPBB-*as conditions for the *[SPB2-, SPBB+]* rule does not change the wild-type phenotype, but leads to anaphase arrest with the decoupled rules. Conversely, deadlocks that appear after the splitting of the *[CK]* rule could be relieved by adjusting the conditions for each variable. Overall, these results underline the importance of variable coupling in the dynamics of this system (and likely of many others), and fully justify using here a partially synchronous formalization.

## Conclusion

Contribution from molecular biology have been essential for our understanding of the mechanisms underlying the cell cycle. So much that, as far as modelling is concerned, they have to a large extent eclipsed the system-level (holistic) view. Indeed, most models of the cell cycle focus on a molecular network built around the cyclin-cdk complexes and their regulators, and forget the other cell components. Paradoxically, cell cycle models do not account for what happens at the cellular level. This was our aim here to take into account all the other components interacting with the “molecular engine” to holistically (and parsimoniously) grasp the whole-cell functioning. Recently, researchers have taken advantage of the wealth of molecular data provided by the -omics revolution, the current understanding of molecular physics, as well as raw computational power, to propose early so-called whole-cell models, which provide realistic simulations of living cells (Karr et al., 2012; Thornburg et al., 2022). However, for the time being at least, the level of details required for such model limits their scope to extremely simple cells, and their complexity hinders their reusability (Waltemath et al., 2016). Even, such attempts appear as a technical feat, instead of providing a clear theoretical synthesis on which to understand the whole cell, as proposed in (Gaucherel, 2019).

Here we developed a complementary and innovative approach. Building on a simple reaction rules formalism, our model provided a basic description of the chromosomal cycle, budding, spindle formation and mitosis of *S. cerevisiae*, explicitly and without resorting to any calculation more complex than Boolean logic. We showed that, in spite of the model’s simplicity, its dynamics faithfully recapitulated the sequence of events of the budding yeast cell cycle. This mainly due to the possibilistic and qualitative properties of such an EDEN formalism (Gaucherel and Pommereau, 2019; Pommereau et al., 2022b).

We used this model to address a concrete biological question, namely the existence of a G2 phase, vs a prometaphase, in *S. cerevisiae*. We showed that our model supports the second proposition, as the G2 phase appears optional, with several pathways directly leading from phase S into metaphase. In contrast, when a prometaphase is considered, all phases follow each other, one after the other, without shortcuts.

From a modelling point of view, our results also support the relevance of synchronous transitions in Boolean models. Similar to the synchronous classes used in GINsim (Fauré et al., 2006; Naldi et al., 2018), reaction rules that affect several variables at once have a strong impact on the dynamics. This suggest that they should indeed be considered as part of the model, and taken into account when analysing the structure of the underlying interaction (regulatory) network.

## Acknowledgements

Preprint version xxx[change to the correct number] of this article has been peer-reviewed and recommended by Peer Community In Mathematical and Computational Biology. (https://doi.org/https://doi.org/10.24072/pci.mcb.101054; A. Fauré, 2026). We wish to thank Franck Pommereau for the EDEN-ecco platform conception and Rudy Usach for early discussions on the subject of this paper.

## Data, script, code, and supplementary information availability

Scripts and code are available online: https://ecco.ibisc.univ-evry.fr/notebooks.html which provides the engine as well as the (Jupyter platform) interface of the EDEN framework. No data were used in this study; the method sources are archived at https://codeberg.org/fpom/ecco.

